# MicroRNA-4776-5p acts as a radiosensitizer and predicts the prognosis of patients with head and neck cancer receiving radiotherapy

**DOI:** 10.1101/2022.12.27.522054

**Authors:** Yo-Liang Lai, Chun-Chieh Wang, Yung-Lun Lin, Pei-Chun Shen, Meng-Hsin Tsai, Fang-Hsin Chen, Wei-Chung Cheng

## Abstract

Head and neck cancer is the leading cancer worldwide. Radiation therapy plays important role of treatment for head and neck cancer. MicroRNAs have been shown to be related to tumor progression and radiosensitivity. However, the mechanisms are still largely unknown and evidence are still limited. In the current study, we sought to identify the miRNA related the radiosensitivity of head and neck tumor cell, which leading to the disappointed prognosis of patients with head and neck cancer receiving radiation therapy. The miRNA expression profiles and clinical information of patients with head and neck cancer were obtained from The Cancer Genome Atlas. The identification of miRNA was carried out through an integrated bioinformatics analysis. The miRNA identified in previous approach was validated through *in vitro* and *in vivo* studies. MiR-4776-5p was finally identified as the role of radio-sensitizer and predicts the prognosis of patients with head and neck cancer receiving radiotherapy. 11 of 16 genes targeted by the miR-4776-5p have been discovered to regulate the mechanisms related to radiosensitivity using functional annotation.

## 1. Introduction

The cancer arised from the mucosal lining of the oral cavity, pharynx, and larynx, are collectively as head and neck cancer (HNC). HNC is the 7^th^ most common cancer worldwide, with more than 660,000 cases and 320,000 deaths per year globally[1]. Among the HNCs, 90% are squamous cell carcinoma (HNSC) whereas the rest are adenocarcinomas. The current standard treatments applied on HNSC include surgery, radiotherapy (RT), chemotherapy, targeted therapy, and immunotherapy. The outcome is improved but still unsatisfied. RT can play a role as either definitive treatment or adjuvant therapy. The postoperative adjuvant radiotherapy has been proved to increase the 5-year cancer-specific survival and overall survival for patients with lymph node-positive HNSC compared with surgery alone [2]. Patients with specific risk factors, such as extracapsular extension (ECE), perineural infiltration, vascular embolism, and lymphatic invasion, are suggested for postoperative radiotherapy or concurrent postoperative radiation plus chemotherapy to achieve better outcomes [3]. However, there is still a group of patients whose disease are prone to progression after this type of intensive treatment. Treatment failure after radiotherapy is correlated to the radioresistance of tumor cells. Therefore, a biomarker used to select those patients with poor response to RT and be a candidate as radiosensitizer are needed.

MicroRNAs(miRNAs) are endogenous RNA with lengths around 19–23 nucleotides that play important regulatory roles by targeting mRNAs for post-translational modifications. miRNAs effectively repress the multiple target genes through imperfect sequence bases, with complementarity, between miRNA and its targets [4]. Abnormal expression of miRNA is found in various tumors which contributes to tumor development, tumor progression, and events leading to treatment resistance [5]. The deregulation of miRNAs in tumors can be harnessed as potential therapeutics by either miRNA replacement therapy using miRNA mimics or inhibition of miRNA function by antimiRs [6]. For example, lipid nanoparticle encapsulated miR-34 mimics has promising anti-tumor activity when applied in animal tumor models of prostate [7], liver [8], and lung [9], and is currently being tested in phase I clinical trial (NCT01829971) in several solid and hematological malignancies [6].

MiRNA modulating the tumor response to ionizing radiation has been reported in several studies [10, 11]. Growing evidences focused on the HNC only [12]. However, most of the miRNAs are identified using microarray methods and less predict the prognosis of clinical patients. Few studies identified the miRNA using high-throughput screening with large sample sizes but lack of evidence performed on *in vitro* and *in vivo* studies [13, 14]. In the present study, we identified a miRNA related to the clinical outcome of patients with HNSC receiving RT alone from The Cancer Genome Atlas (TCGA) database, and validated it on *in vitro* and *in vivo* experiments. This miRNA could be potentially selected as miRNA-based therapeutic targets to block the radioresistant characters and lead to better treatment outcomes for head and neck cancer.

## 2. Results

### 2.1. Identify the clinically radiosensitive MIRNA of patients with head anc neck cancer

In order to identify the candidate miRNA acting as radiosensitizer for head and neck cancer cell, we conducted an integrative study combining the clinical cohort and bench data (Figure 1). Firstly, The Cancer Genome Atlas Head-Neck Squamous Cell Carcinoma (TCGA-HNSC) data with the total of 528 patients were enrolled. Only 432 patients have the record of confirmatory receiving RT status, including “yes” with 294 patients and “no” with 138 patients (Table1 & supplemental Table I). 294 of 432 patients have received radiotherapy. In order to focus on the role of “radio-sensitizer” only, the patients who have received any other systemic therapy (chemotherapy, targeted therapy…) were excluded. Finally, 128 patients received radiotherapy alone. Then a total of 2588 miRNAs were used as the possible candidate. After differential expression (DE) analysis, 277 miRNAs showed significant difference between tumor and adjacent normal tissue in these patients. Using the overall survival (OS) and progression free interval (PFI) as end-points of survival analysis, 12 miRNAs were determined to be significantly related to the disease progression of the patients with HNSC receiving radiotherapy alone. Hazard ratio (HR) <1 was used to select the miRNA related to better prognosis (radiosensitizer) and the miRNAs which also meet the above criteria in the subgroup of patients treating without RT were excluded. Finally, miR-4776-5p was identified as the candidate for further functional annotation and validation experiment. The Kaplan-Meier survival plotter showed the expression of mir-4776-5p are significantly related to clinical prognosis in HNSC-RT(+) group but not in HNSC-RT(-) group, which might indicate the prognostic role of mir-4776-5p contributed by radiosensitivy rather than tumor behavior (Figure 2).

**Figure 1:**
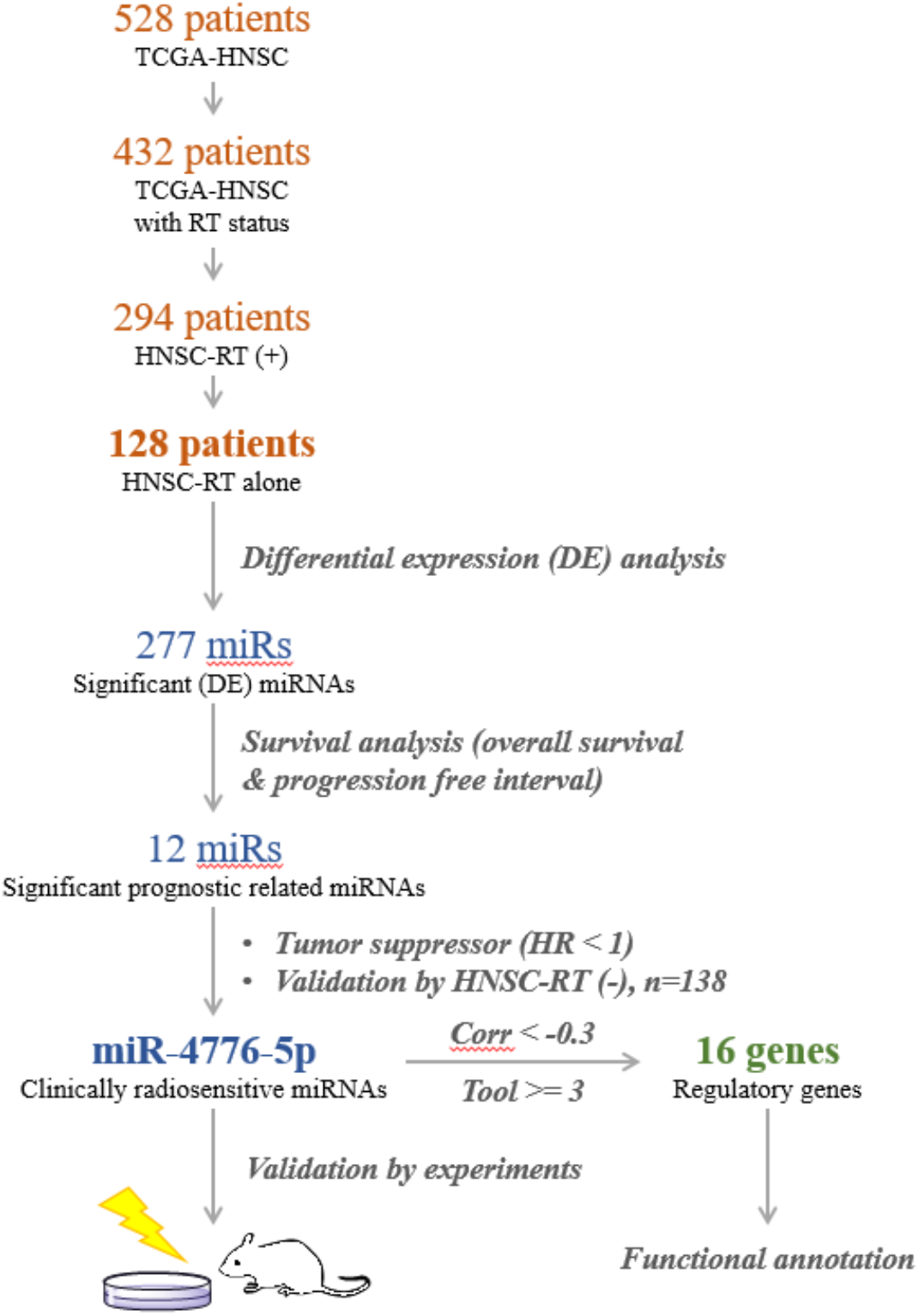
pipeline of the study

**Table 1:** Basic characteristic of patient receiving RT alone

| Characteristic | RT alone (N = 128) |
| --- | --- |
| **Age at diagnosis—yr** |  |
| Median (IQR) | 64 (55-70) |
| ≥65 (%) | 62 (48.4) |
| <65 (%) | 66 (51.6) |
| **Sex —no. (%)** |  |
| Male | 92 (71.9) |
| Female | 36 (28.1) |
| **Stage —no. (%)** |  |
| I | 5 (4.1) |
| II | 14 (11.4) |
| III | 31 (25.2) |
| IV | 73 (59.3) |
| **Neoplasm histologic grade —no. (%)** |  |
| GX | 2 (1.6) |
| G1 | 17 (13.3) |
| G2 | 72 (56.2) |
| G3 | 37 (28.9) |
| **Smoking** |  |
| Yes | 100 (80.0) |
| No | 25 (20.0) |
| **Alcohol** |  |
| Yes | 90 (72.6) |
| No | 34 (27.4) |

**Figure 2:**
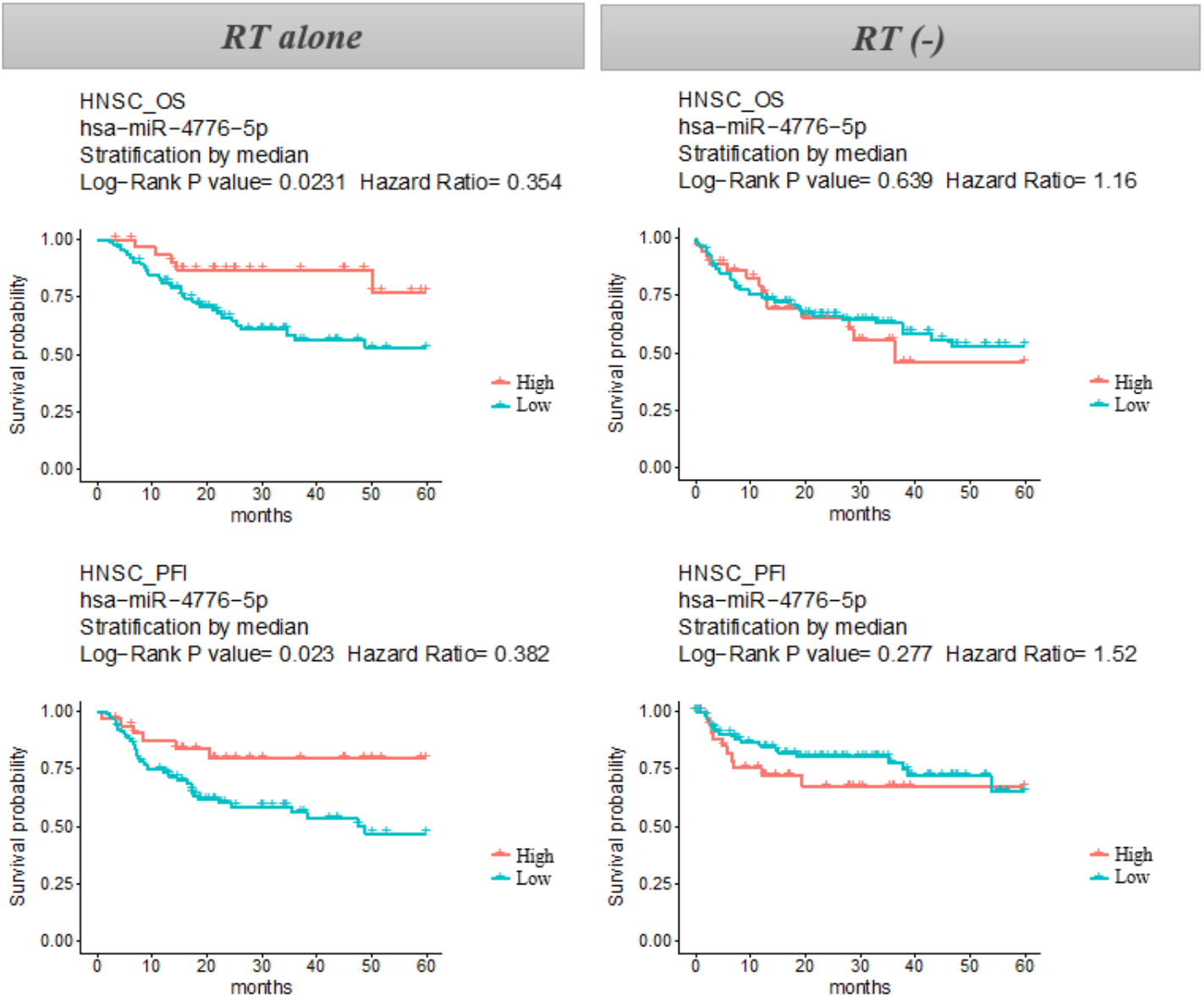
Kaplan-Meier survival plotter of overall survival (OS) and progression free interval (PFI) in patients with head and neck cancer receiving RT alone and without RT.

### 2.2 Identify the target gene and functions regulated by MIR-4776-5p

To investigate the mechanism of miR-4776-56 acting as a radiosensitizer, we identified the target genes of miR-4776-5p and their functions in a systematic approach (see Identification of targeted gene and functional annotation in Material and Methods for details). Briefly speaking, the targeted genes had negative correlation with miRNAs in expression level. The results had to be also confirmed by at least 3 target prediction tools, which has been used in our previous publications[15, 16]. 16 genes (A*P1M1, ATAD3B, CLEC11A, CMTM3, COLGALT1, FBX044, KDELR1, MCTS1, MRPL17, MTCH1, PARVB, PLOD1, PRAF2, RPS27L, SH2D2A, VKORC1*) have been identified and the functional annotation was performed (supplemental table 2). The functions related to radiosensitivity, including immune system, mitochondria, signal transduction, apoptosis, VEGF pathways, cell proliferation, DNA damage response, and hypoxia, were summarized with the regulatory network with targeted genes in Figure 3.

**Figure 3:**
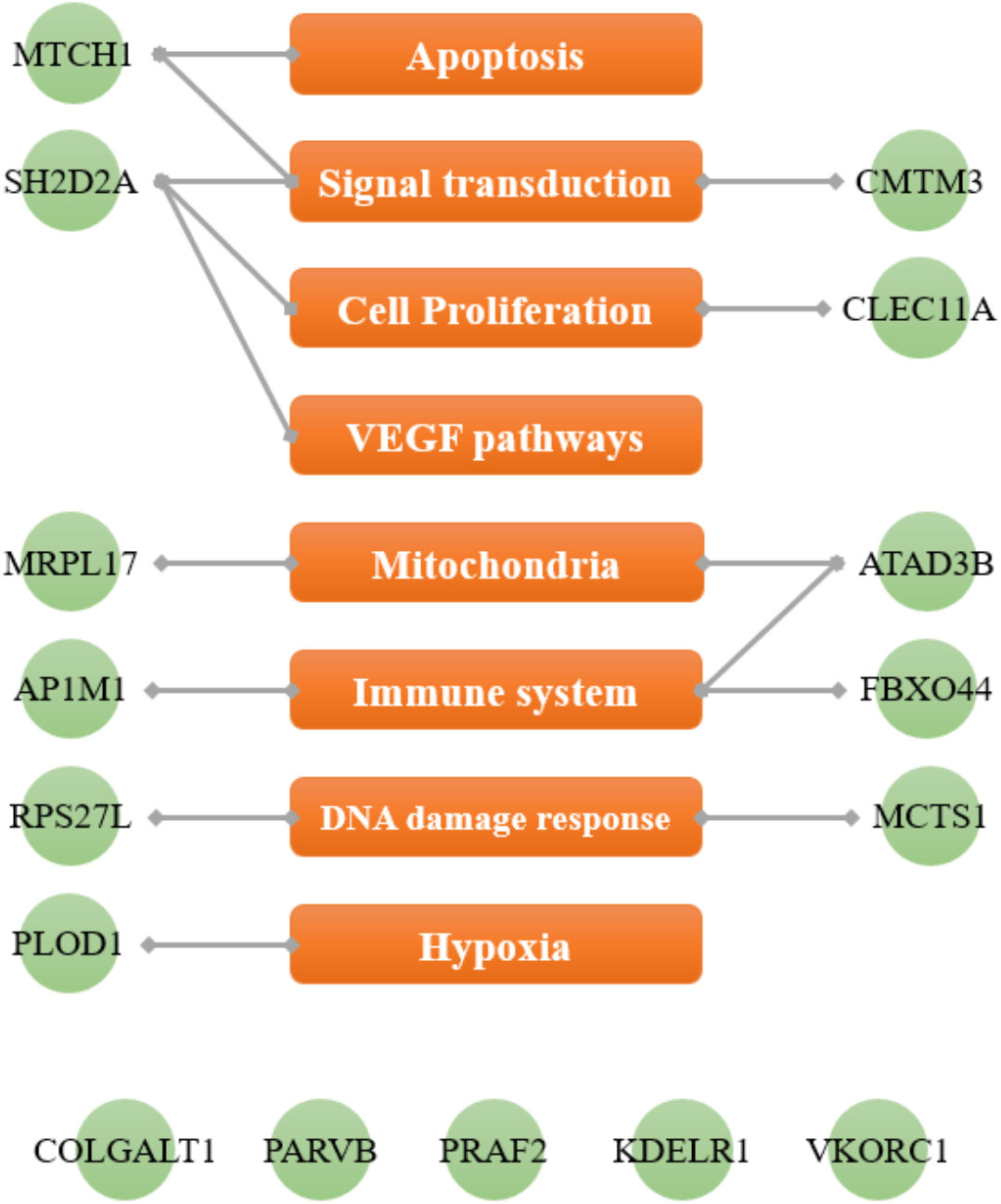
The targeted genes of miR-4776-5p and the possible regulatory functions related to radiosensitivity.

### 2.3 MiR-4776-5p suppress DNA repair and sensitizes HNSC cells to ionizing radiation

To explore the effect of miR-4776-5p on DNA damage and repair efficiency of Fadu cells, the extent of DNA double strand break (DSB) indueced by RT was examined. The phosphorylation of H2AX is a well-known marker to assess the formation of DSB and the following repairing kinetics. We therefore quantified the number of γH2AX foci at different time points after RT in cells trasfected with miR-4776-5p or NC. As indicated in Figure 4, 2Gy RT led to a remarkable increasing γH2AX foci 30 min post-RT either in Fadu cells transfected with miR-4776 mimic or NC miRNA. The expression of miR-4776-5p persisted the γH2AX foci number until 8 hr post-RT comparing with that of NC miRNA, indicating the un-efficient DNA repairing machinery. In addition, a clonogenic assay was performed following transfection of Fadu cells with miR-4776-5p and NC miRNA to evaluate the effect of miR-4776-5p overexpression on the radiosensitivity of Fadu cells. Results of the survival curve showed significantly moved down with the increase of doses in cells transfected with miR-4776-5p compared with those transfected with NC miRNA (p < 0.001). These results suggested that miR-4776-5p enhanced the radiosensitivity of Fadu cells in a manner that may be involved in the persistence of DNA repair and suppression of cell viability

**Figure 4:**
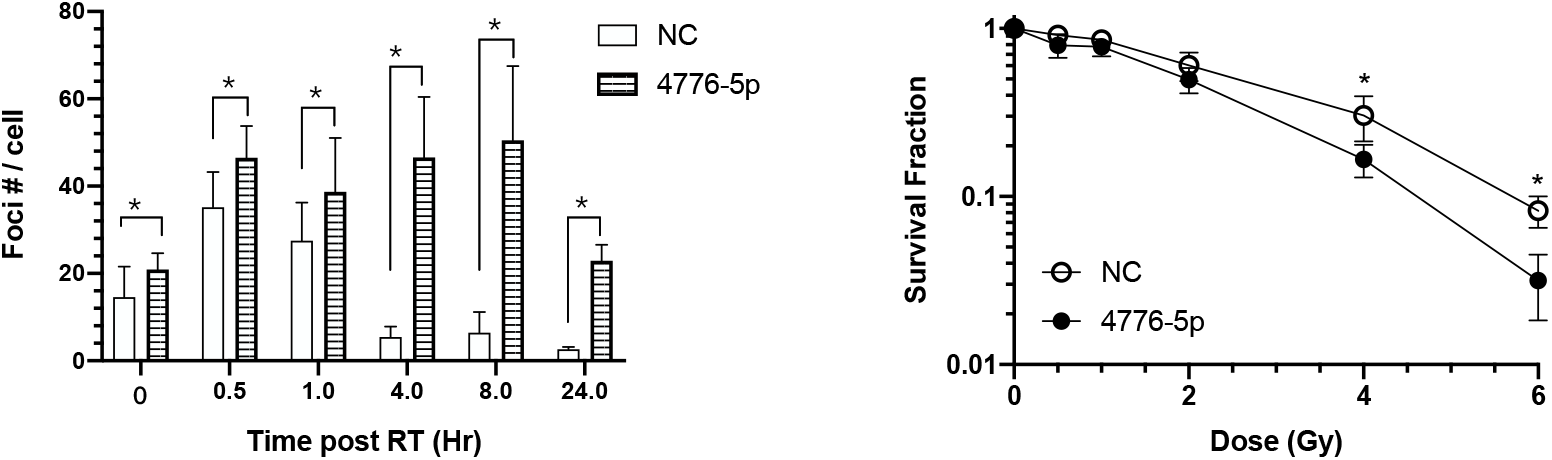
Expression of high mir-4776 expression on the radiosensitivity of Fadu cells. (A) Quantification of the DNA double strand breaks levels by γH2AX foci numbers in cells with or without overexpression of miR-4776-5p (B) Analysis of surviving fractions of Fadu cells exposed to gradient doses of Cs-137 with or without overexpression of miR-4776-5p. *: *p* < 0.001

### 2.4 Tumor growth in response to over-expression of miR-4776-5p and ionizing radiation

Increasing the level of miR-4776-5p in Fadu cells had significant effect on suppressing DNA double strand break repair and inducing mitotic death. We therefore investigate this in an in vivo system to study whether increase of radiosensitivity is relevant on tumor growth after ionizing radiation. Specially, Fadu cells were transfected with miR-4776-5p mimic or scramble NC for 24 hr, and then inoculated into immune-deficient BALB/C nude mice. Mice bearing Fadu-NC or Fadu-miR-4776-5p tumors were treated by 13Gy RT when tumor volume reached 150mm^*3*^. Transfection of miR-4776-5p did not affect tumor growth rate of Fadu since it took 9 days for tumor grew up to 600mm^*3*^, as same as the NC group. Treatment with ionizing radiation retarded the growth in tumors of NC which took 17 days to reach 600mm^*3*^. On the other hand, ionizing radiation also greatly affect tumor growth on miR-4776-5p transfected tumors which took 19 days to reach 600mm3 and remain the same tumor size up to 29 days. Comparing to the non-RT group, the average tumor growth delay trigger by RT is 8 days for NC, and 20 days for mir4776 group (Figure 5). This data again suggest that miR-4776-5p overexpression increase to radiosensitivity of HNSCC cells.

**Figure 5:**
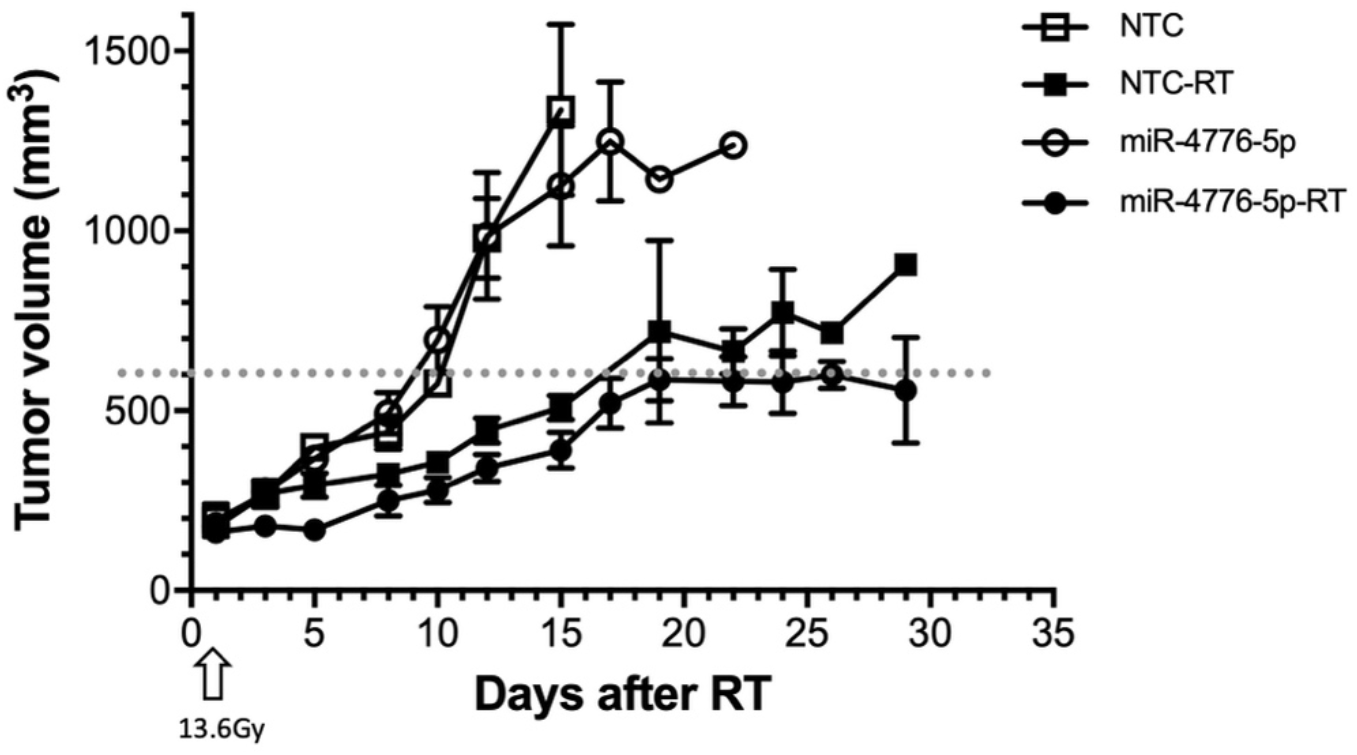
Mir-4776-5p enhanced the tumor suppression capability of RT in vivo. Fadu/miR-NC or Fadu/mir-4776-5p cells were subcutaneously inoculated into SCID mice. the mice were exposed to RT at a dose of 13.5 Gy when tumor volume reached 150mm^3^. Tumor volumes were measured every 2 days after RT.

## 3. Discussion

HNC occurs as one of the most common 10 cancers worldwide, with the 5-year survival being about 40% even if receiving anti-cancer therapy [1, 17]. RT is important part of the definitive or adjuvant treatment of HNC. Resistance to RT, so called radioresistance, plays important role of treatment failure of HNC. MiRNAs include a group of smRNAs with 19-23 nucleotides long, regulating the growth of tumor cell. To date, several miRNAs that have shown be related to the diagnostic, prognostic, or therapeutic in HNC have been reported[18]. Several miRNAs have also shown the efficacy of regulating tumor radiosensitivity by modulating several pathways[19]. However, the results from above literatures are not consistent between bench and clinical condition, limiting the usefulness in precision medicine era. Therefore, we conduct current study to investigate the candidate miRNA which not only regulate the tumor radiosensitivity *in vtiro* and *in vivo* studies but also predict the prognosis of patients with HNC receiving RT. In our study, we identified miR-4776-5p as a prognostic biomarker from the clinical cohort including patients with HNC receiving RT alone through an integrated bioinformatics approach, and validate the regulation of radiosensitivity of it in both *in vitro* and *in vivo* studies using head and neck tumor cell. This miRNA has been reported in miRNA-gene regulatory network in gastric cancer only previously[20]. The consistent result in our study suggested miR-4776-5p have strong potential to be used to develop a novel radiosensitizer to improve the treatment outcome in HNC.

To understand the pathways regulating by miR-4775-5p, we first identified the target genes of the miRNA using miRTarBase database and several bioinformatics tools for miRNA target prediction. 16 genes, including *(AP1M1, ATAD3B, CLEC11A, CMTM3, COLGALT1, FBX044, KDELR1, MCTS1, MRPL17, MTCH1, PARVB, PLOD1, PRAF2, RPS27L, SH2D2A, VKORC1)*, have been identified. AP1M1, which is an intrinsic part of the clathrin adaptor AP-1 complex. Previous studies suggested that *AP1M1* involved in the progression of liver cancer caused by HBV infection[21] and identified as biomarker related to central nervous system (CNS) metastasis of breast cancer[22]. ATAD3B is a mitochondrial membrane-bound ATPase belongs to ATPase family AAA domain-containing protein 3 *(ATAD3)*, which linked to the progression and prognosis of various cancer types including liver cancer[23], breast cancer[24], and CNS cancer[25]. A global proteome analysis of the mitochondria in Raje cell also showed ATAD3B as the potential biomarkers of radioresistance[26]. *CLEC11A* is a protein coding gene significantly participating in tumor microenvironment and immune cells’ communication, and can predict the survival of protein in laryngeal squamous cell carcinoma (LSCC) [27]. *CMTM3* is not only related to the cell growth and migration of oral squamous cell carcinoma [28] and pancreatic cancer [29], but also has the role in tumor microenvironment and cancer immunotherapy [30]. *COLGALT1* has been reported as a biomarker for predicting prognosis and immune responses for kidney renal clear cell carcinoma [31]. *KDELR1* involved the regulation of T cell homeostatsis [32] and correlated with the prognosis glioma [33]. *MCTS1* has been identified as oncogenes in various tumor entities including HNSC [34] and correlated to immune cell infiltration [35]. *MRPL17* has been implicated as crucial player in oral carcinogenesis from analysis in patients with tongue cancer [36]. Upregulation of *MTCH1* is associated with cell proliferation of liver hepatocellular carcinoma [37]. The increasing presentation of *MTCH1* neoantigen after ionizing radiation was also noted [38]. *PARVB* has been suggested as a clinically useful biomarker for adjuvant tongue squamous cell carcinoma based on the function of increasing cell migration capability [39]. *PLOD1*, which belongs to the Procollagen-Lysine, 2-Oxoglutarate 5-Dioxygenase (PLOD) gene family, which is related to multiple cancer entities, has shown to play the significant role in the development of LSCC [40]. *PRAF2* is involved in the occurrence and progression of several malignant tumors, including esophageal, liver and brain cancer [41-43]. *RPS27*, which encodes MPS-1 protein, has been reported to be involved DNA repair and regulate radiation sensitivity via the MDM2-p53 and MDM2-MRN-ATM axes [44, 45]. *SH2D2A*, as a gene encoding an adaptor protein thought to function in T-cell signal transduction, has been identified as radiation response gene in a study irradiating primary human fibroblast cell lines [46].

To determine the mechanisms of miR-4776-5p acting as a radiosensitizer, we performed functional annotations of the 16 genes targeted by miR-4776-5p. The result pointed out that the cross-reaction of 11 among 16 genes were involved in the functions including DNA damage response, apoptosis, cell proliferation, signal transduction, hypoxia, VEGF pathways, mitochondria, immune system. DNA damage response and apoptosis were recognized as the traditional mechanism of radiation inhibiting tumor cell proliferation[47, 48]. Increasing evidence presented the functional links between signal transduction and cellular responses [49]. Hypoxia is a common feature of the microenvironment in tumors and associated with the resistance to RT [50]. VEGF knockdown has shown to enhance radiosensitivity by inhibiting the occurrence of autophagy [51]. Cellular organelles, in particular mitochondria, has been noted to mediate the radiation response in tumor, as regulating many of the cellular processes involved in radioresistance [52]. Recent data suggested that radiation not only kill the local tumor but also inhibit the growth of distant metastatic lesions with immunomodulatory properties (so called abscopal effect) [53, 54]. To sum up, these functions conformed the feasibility of miR-4776-5p act as a candidate regulating radiosensitivity and the potential for further radiobiological research.

Although there have been numerous studies investigating the role of miRNAs in modulate with radiosensitivity of HNC, few of them presents the consistent results between bench and clinical condition [55]. In most of previous studies, researchers investigate the expression change of miRNAs from HNC cell lines *in vitro* using microarray detection first to identify candidate miRNA. Few of them then validated the candidate miRs in clinical samples [56, 57] Recently, with the revolution of high throughput Next Generation Sequencing (NGS) methods (which has higher sensitivity than micro-array), several large sequencing database including TCGA have been conducted with big amount of sequencing data and large clinical sample size. To the best of our knowledge, only two studies has investigated miRNA related to radio-sensitivity of patients with HNC from these big data with integrative bioinformatics approaches[13, 14]. Using TCGA data, a 5-microRNA signature-based nomogram has been proposed to predict the response to radiotherapy of HNSC patients *in silico* data analysis only [13]. Inoue et al [14] also using TCGA data for screening first but focused on human papillomavirus (HPV)-negative oropharyngeal squamous cell carcinoma (OPSC) only and suggested miR-130b has potential act as a biomarker for the radiosensitivity of HPV-negative OPSC. However, there are some limitations in these two studies. First, the patient receiving RT may also receive chemotherapy or targeted therapy. It’s is difficult to determine the change of prognosis contributed by which type of treatment. Second, the result has not been validated in bench studies to confirm the result is related to radiosensitivity. Therefore, in our study, we screen the miRNAs from the patient receiving RT **alone**, and validate the *in silico* result with the radiobiological studies *in vitro* and *in vivo*. Besides, we also validated our candidate miRNA in the clinical cohort of HNSC receiving **no** RT. The inconsistence result between group of RT alone and no RT favored that miR-4776-5p regulate the radiosensitivity and response of radiotherapy instead of HNC progression itself.

The results of biological validation of miR-4776-5p showed its great impact on DNA repair efficiency and the survival of HNC cells after irradiation. By quantification of the γH2AX foci number we observed that DSB resolution was greatly interfered in irradiated cells with miR-4776-5p overexpression. Overexpression of miR-4776-5p triggered a higher number of foci at 2.5 h after RT than cells treated with miR-NC and showed inefficient DSB repair kinetics within the 24 h interval after RT. Our result also showed that overexpression of miR-4776-5p had significant impact on the survival of HNC cell after RT due to the unrepaired DNA aberrations and failed cell division. In vivo study showed that overexpression of miR-4776-5p did not affect the tumorigenesis of HNSC cells, but enhanced tumor radiosensitivity, regarding of the prolonged tumor growth inhibition in tumor cells transfected with miR-4776-5p.

## 4. Materials and Methods

### 4.1 Data collection and processing

The smRNA and RNA sequencing profiles and clinical information of TCGA-HNSC dataset were obtained from our previous studies: YM 500 and Driver DB database. RNA-seq and clinical data, such as the primary tumor as well as normal and metastatic tissues, are retrieved and annotated from the TCGA data portal (https://portal.gdc.cancer.gov/). As to the smRNA-seq data, it is downloaded from CGHub (https://cghub.ucsc.edu/) and then processed by the miR-seq pipeline of YM500 [58] and annotated with miRBase database R21[59] and DASHR database v1.0[60]. This study curates 20,495 genes and 2588 miRNAs in total for analysis.

### 4.2 DE analysis and survival analysis

Differentially expressed analysis aims to define significant differentially expressed miRNAs between primary tumor samples and adjacent normal samples. For this purpose, an R package, DEseq (Version 1.28.0)[58], is applied to filter candidates based on the criteria of adjusted p-values < 0.05 and log2 fold-change values greater than 3 for miRNAs and efficiently identify oncogenic expression between tumor samples and normal samples. Furthermore, candidates with normalized mean counts (baseMean) < 1 for miRNAs are filtered out to eliminate candidates with excessively low expression levels. In the survival analysis, the aim is to define clinically relevant candidates. Survival (Version 2.41–3), an R package, is applied to calculate the Cox proportional hazards model between two predefined groups, providing survival estimates for time-to-event datasets.

### 4.3 Identification of targeted gene and functional annotation

To confirm the relationships between miRNAs and target genes, we performed the following two analytical steps. The first step is calculating correlations between gene and miRNA expression levels by Pearson’s, Spearman’s, and Kendall’s coefficients. This method comprises both linear and non-linear correlation, which enhances the sensitivity of detecting potential target genes of miRNA candidates. Genes and miRNAs which any of the three correlation coefficients < − 0.3 are considered. The second step is investigating the interactions between miRNAs and their target genes by twelve additional bioinformatic prediction tools, which predicted the interaction between miRNA and corresponding target genes. As detailed in the previous publications[58], only the genes satisfied the following criteria: 1) correlation coefficient was <− 0.3 in first step, 2) the interaction was supported by more than 3 bioinformatic tools, was identified as the targeted gene. Functional annotation of the targeted gene was performed using GeneCards database[58].

### 4.4 In vitro study

#### 4.4.1 Cell culture, miRNA transfection and irradiation

Fadu cells (HTB-43TM, ATCC) were maintained in MEM medium supplemented with 10% fetal bovine serum, 1% antibiotics and maintained at 37°C in a humidified atmosphere containing 5% CO2 condition. Cells were transfected with selected 4776-5p mimics and Negative Control (mirVana miRNA Mimics, Applied Biosystems, Thermo Fisher Scientific) at a final concentration of 40nM using Lipofectamine 3000 (Invitrogen) according to the manufacture’s instruction and keep maintenance to the study time points. Irradiation was delivered by 6-MV X-ray from a linear accelerator (Clinac® iX, Varian Medical System) with a dose rate of 6 Gy/min).

#### 4.4.2 Quantitative real-time PCR

After transfected with synthetic mimics, cells were collected at 24 hr in Trizol reagent to extract the total RNA component. MicroRNA was firstly transcribed by miRCURY LNA RT kit to obtain its cDNA and examined for the expression level by specific primers designed for miR-4776-5p (Applied Biosystems). The quantitative PCR was performed by the miRCURY LNA SYBR Green PCR KiT (Applied Biosystems) and analyzed by CFX ConnectTM Real-Time PCR Detection System (Bio-Rad, Hercules, CA, USA). The relative expression level of each miRNA was compared to the RNU6 endogenous control and normalized to cells transfected with scramble miRNA (NC group) using the 2 − ΔΔCt method.

#### 4.4.3 Cell viability after RT

Fadu cell transfected with synthetic mimics were irradiated with different doses to test its intrinsic response to RT. Briefly, Fadu cells were plated in each well of 96-well plates at a density of 5000 cells per well in 200 μL of culture medium and maintained at 37 °C. After 24hr, cells transfected with synthetic mimics and incubated for 24hr. Cells were irradiated with 2 and 4Gy and developed using Cell Counting kit -8, CCK-8 (Biotools, Taiwan) at day3 after RT. The amount of the formazan dye in survived cells was determined by measuring O.D. at 450nm from five independent experiments which was directly proportional to the number of living cells.

#### 4.4.4 Colony formation assay

The transfected Fadu cells were trypsinized and suspended in 1X PBS, and then exposed to radiation (0-6Gy). Cells were pated in 6-well dishes at different densities depending on the potency of the irradiation treatment (from 400 to 3200 cells/well). Ten days later, the cells were fixed stained with crystal violet in 70% ethanol. The number of colonies, defied as >50cells/colony were counted. The survival fraction was determined as the ratio of the number of colonies in the treated groups to the number of colonies in the untreated group. Six wells were set up for each condition.

#### 4.4.5 Immunohistochemistry staining

For immunostaining of γH2AX, after 30minutes, 1 hour, 4 hours, 8 hours and 24hours of radiation exposure, Fadu cells were fixed for 15 minutes at room temperature with 4% paraformaldehyde and washed twice with PBST (1x PBS with 0.1% Tween-20). After washing, cells were incubated with Permeabilization buffer (0.3% Triton X-100 in PBS) for 10 minutes at room temperature then with blocking buffer (3% BSA in PBS) for one hour at room temperature. After washing, slides were incubated with primary rabbit anti-γH2AX antibody (Merck) diluted in blocking buffer at 4 °C overnight and then incubated for 30 minutes at room temperature with anti-rabbit IgG coupled with FITC (BD). After washing, slides were mounted with mounting medium containing DAPI (Life Technologies) for nucleus staining. Images were automatically acquired by ImageXpress Micro Confocal system (Molecular Devices, San Jose, CA, USA). At least 250 Nuclei were captured and the number of γH2AX foci/nuclei was analyzed using MetaXpress software (Molecular Devices, San Jose, CA, USA)

### 4.5 In vivo study

To examine the effect of miRNA interfering tumor response to radiotherapy, 8-week-old male mice (BALB/c nu/nu, National Science Council Animal Center, Taipei, Taiqan) were subcutaneously injected with 3 X10^6^ viable transfected Fadu cells, in a total volume of 100 uL of PBS. After implantation, tumor volume of the xenografts was measured using caliper according to the formula π/6 × (large diameter) × (small diameter)2. Tumor-bearing mice received local irradiation for 13Gy as tumor volume reached 150mm^3^. To access tumor response to irradiation, tumor volume was continuously measured to determine the growth rate until reaching 1200 mm^3^. The protocol of irradiation was described in details in our previous publication [59]. Briefly, tumor-bearing mice were anesthetized by a mixture (1:1) of ketamine (50 mg/kg, Merial Laboratoire de Toulouse, Toulouse, France) and xylazine (20 mg/kg, Bayer HealthCare Animal Health, Leverkusen, NRW, Germany) and were restrained during irradiation. The tumors were covered with a 1-cm bolus on the surface and irradiated by 6 MV X-rays from a linear accelerator with a dose rate of 2–3 Gy/min. All animal experiments were performed with the approval of the Institutional Animal Care and use Committee (IACUC) of Chang Gung Memorial Hospital (CGU110-071)

### 4.6 Statistical analysis of invivo and invitro study

Statistical analyses for biological experiment were performed using the Student’s t-test for a simple comparison of the two groups. Differences were considered statistically significant if the P value was < 0.05

---

**Supplemental table 1:** Basic characteristic of patient receiving no RT.

| Characteristic | RT (-), N = 138 |
| --- | --- |
| **Age at diagnosis —yr** |  |
| Median (IQR) | 63 (58-75) |
| ≥65 (%) | 61 (44.5) |
| <65 (%) | 76 (55.5) |
| **Sex —no. (%)** |  |
| Male | 87 (63.0) |
| Female | 51 (37.0) |
| **Stage —no. (%)** |  |
| I | 12 (8.9) |
| II | 47 (35.1) |
| III | 28 (20.9) |
| IV | 47 (35.1) |
| **Neoplasm histologic grade —no. (%)** |  |
| GX | 1 (0.7) |
| G1 | 23 (16.7) |
| G2 | 85 (61.6) |
| G3 | 29 (21.0) |
| **Smoking** |  |
| Yes | 97 (73.5) |
| No | 35 (26.5) |
| **Alcohol** |  |
| Yes | 79 (59.8) |
| No | 53 (40.2) |

**Supplemental table 2:**
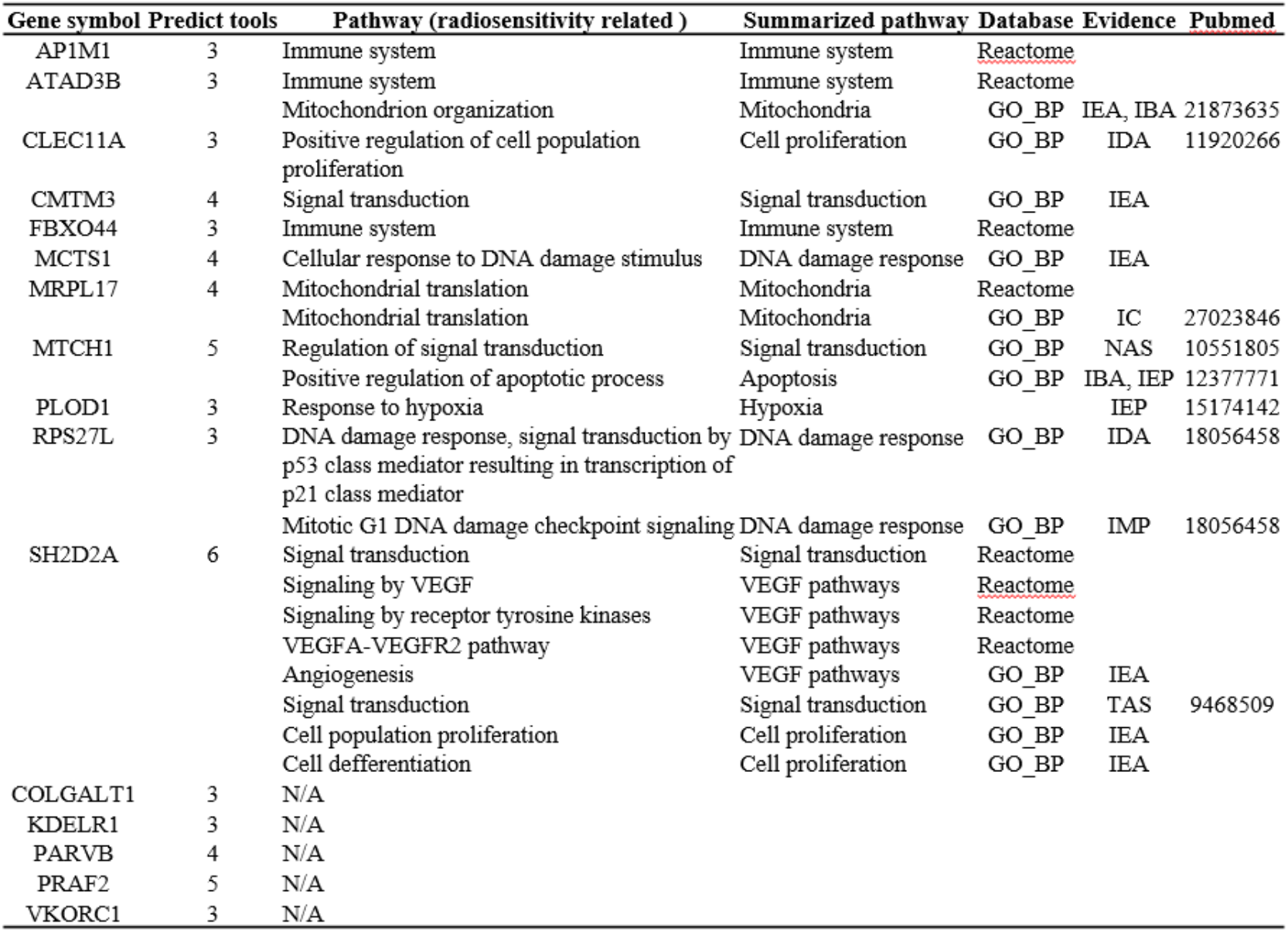
16 targeted genes and functions related to radiosensitivity.

